# LCSkPOA: Enabling banded semi-global partial order alignments via efficient and accurate backbone generation through extended LCSk++

**DOI:** 10.1101/2024.07.18.604181

**Authors:** Minindu Weerakoon, Christopher T Saunders, Haynes Heaton

## Abstract

**Background:** Most multiple sequence alignment and string-graph alignment algorithms focus on global alignment, but many applications exist for semi-global and local string-graph alignment. Long reads require enormous amounts of memory and runtime to fill out large dynamic programming tables. Effective algorithms for finding the backbone and thus defining a band of an alignment such as the longest common subsequence with kmer matches (LCSk++) exist but do not work with graphs. This study introduces an adaptation of the Longest Common Subsequence with kmer matches (LCSk++) algorithm tailored for graph structures, particularly focusing on Partial Order Alignment (POA) graphs. POA graphs, which are directed acyclic graphs, represent multiple sequence alignments and effectively capture the relationships between sequences. Current state of the art methods like ABPOA and SPOA, while improving POA, primarily focus on global alignment and thus are limited in local and semi-global banding scenarios. Our approach addresses these limitations by extending the LCSk++ algorithm to accommodate the complexities of graph-based alignment.

**Results:** Our extended LCSk++ algorithm integrates dynamic programming and graph traversal techniques to detect conserved regions within POA graphs, termed the LCSk++ backbone. This backbone enables precise banding of the POA matrix for local and semi-global alignment, significantly enhancing the construction of consensus sequences. Compared to unbanded semi-global POA, our method demonstrates substantial memory savings (up to 98%) and significant run-time reductions (up to 37-fold), particularly for long sequences. The method maintains high alignment scores and proves effective across various string lengths and datasets, including synthetic and PacBio HiFi reads. Parallel processing further enhances runtime efficiency, achieving up to 150x speed improvements on conventional PCs.

**Conclusion:** The extended LCSk++ algorithm for graph structures offers a substantial advancement in sequence alignment technology. It effectively reduces memory consumption and optimizes run times without compromising alignment quality, thus providing a robust solution for local and semi-global alignment in POA graphs. This method enhances the utility of POA in critical applications such as multiple sequence alignment for phylogeny construction and graph-based reference alignment.

## Background

Graph-based aligners, such as GraphAligner[1], have become increasingly important in genomics and bioinformatics due to their ability to provide a more flexible and comprehensive representation of genetic variation compared to linear sequence-based reference mapping techniques like minimap2[2], HISAT2[3], and BWA[4]. These graph-based reference alignment techniques decrease reference bias and enhance the accuracy of variant detection and genome assembly. Graph-based reference alignment, despite its advantages, is much slower than linear sequence-based methods. Since the creation of graph-based references depends on multiple sequence alignment, speeding up and improving multiple sequence alignment will help reduce the performance gap. While multiple sequence alignment is a well-established problem with many publications addressing and improving it, challenges still exist.

This study introduces a novel adaptation of the Longest Common Subsequence with k-mer matches (LCSk++) algorithm specifically designed for graph structures, with a focus on enabling banded local and semi-global Partial Order Alignment (POA) graphs [5]. POA graphs are directed acyclic graphs that represent multiple sequence alignments, capturing the relationships between sequences and accommodating variations such as insertions, deletions, and substitutions more flexibly and comprehensively than traditional alignment methods.

Existing methods like ABPOA [6] and SPOA [7] improve POA by using greedy banding and computing the full matrix with use of SIMD optimizations, respectively. However, these methods are limited and cannot effectively apply to local and semi-global alignment, which are crucial for many alignment use cases. Our approach extends the LCSk++ algorithm [8, 9] to address these complexities, enabling efficient banded semi-global POA.

Our approach extends the LCSk++ algorithm, originally developed for linear string sequences, to handle the intricacies of graph data structures. By integrating dynamic programming and graph traversal techniques, we enable the detection of conserved regions within POA graphs. This conserved region, which we call the LCSk++ backbone, allows us to band the POA matrix to accurately locally and semi-globally align sequences. This method significantly enhances the construction of consensus sequences with high accuracy and efficiency. Additionally, it allows for other critical applications of POA, such as multiple sequence alignment (MSA) for phylogeny construction and graph-based reference alignment.

Significantly, our method achieves a 37-fold increase in speed and a 98% reduction in memory usage for sufficiently long sequences. Additionally, we can achieve up to 150x speed improvements on conventional PCs through parallel processing without compromising alignment quality, a feat that is not possible with unbanded POA. This advancement allows for fast and efficient local and semi-global alignment on the banded POA graph, enhancing its utility in bioinformatics.

Overall, the extended LCSk++ algorithm for graph structures represents a significant step forward in sequence alignment technology, offering improved performance and accuracy for the analysis of complex biological datasets.

### Preliminaries

In this section we formalize all the required concepts and definitions used in the rest of the paper.

#### Sequence alignment in bioinformatics

In bioinformatics, aligning biological sequences—composed of nucleotide bases A (Adenine), C (Cytosine), G (Guanine), and T (Thymine)—is a fundamental task for deriving consensus sequences. Consensus sequences represent the most common nucleotide or amino acid at each position within a set of aligned sequences. These are invaluable for identifying conserved regions critical to protein function and predicting regulatory elements in DNA. When aligning multiple sequences, pairwise alignment using the Smith–Waterman algorithm [8] has a time complexity of O(L^N^), where L is the sequence length and N is the number of sequences. However, this approach becomes computationally infeasible for large numbers of sequences (N > 5). To address this, multiple sequence alignment through partial order alignment (POA) is employed. POA constructs a graph where nodes represent individual nucleotide bases and edges indicate the level of agreement among sequences. The graph is then aligned to a linear sequence, and subsequently updated based on the alignment. This method achieves a linear time complexity of O((2n+1)LMN), where n is the average number of predecessor nodes per node in the POA graph, M is the number of nodes in the graph, N is the number of sequences, and L is the length of the aligning sequence [5]. While the POA algorithm is more memory- and time-efficient than pairwise alignment, it still demands substantial resources when applied to reads from current long-read sequencing technologies, such as PacBio HiFi which contain reads of length >10,000 BP. Employing efficient banding techniques to calculate only the essential regions of the POA matrix can significantly reduce L and M, making the process feasible on consumer-level hardware.

#### LCSk++

LCSk, which stands for Longest Common Subsequence of k-mers, identifies common subsequences which are exactly k-length k-mers in a pair of sequences [9]. LCSk++ builds upon this algorithm by extending the capability to consider k-mers of lengths greater than k [10]. Our algorithm further builds on this approach to address the case of aligning a POA graph to a sequence. The following definitions outline the core concepts of LCSk++, along with additional definitions, which are utilized in our algorithm.

##### Definition 1 (Common subsequence)

Let X and Y be two strings. Define two sets of indices I = {i_1_, i_2_, …, i_n_} and J = {j_1_, j_2_, …, j_n_}, where the indices in each set are ordered such that i_1_ < i_2_ < … < i_n_, j_1_ < j_2_ < … < j_n_. For every x in 1,…,n, the condition X_ix_ = Y_jx_ holds. These index sets define a subsequence shared by X and Y with a length of n.

##### Definition 2 (k++ common subsequence)

Given a common subsequence of strings X and Y, this subsequence defines two distinct sets of indices I and J (as described in Definition 1). If these sets can be divided into groups of consecutive indices, where each group contains at least k indices, then the subsequence is referred to as a k++ common subsequence.

##### Definition 4 (LCSk++).

LCSk++ of two strings X and Y is the length of their k++ common subsequence with maximal number of elements.

##### Definition 5 (LCSk++ graph)

LCSk++graph of a POA graph and string Y is identifying k++ common subsequence between all the paths (sequences) of a POA graph and the sequence Y with the maximal number of elements and finding the path sequence pair with max number of elements.

##### Definition 6

A substring of string X that begins at index i and ends at index j is represented as X_i_…_j_. If i > j the substring X_i_…_j_ is considered an empty string.

### Implementation

#### Basic dynamic programming

Finding the optimal graph to sequence LCSk++ path can be done using basic dynamic programming recursive relation similar to that of [10] with few modifications. In our implementation, we adopt the definitions as described by [12] with some additional definitions to accommodate the graph structure as presented below.

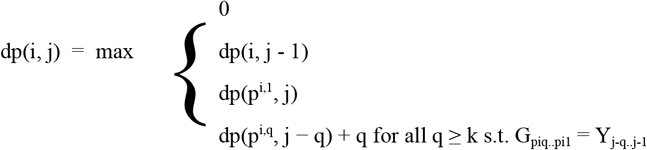

Where,

dp(i, j): entry in dynamic programming tables row i and column j, where i is the i^th^ topologically ordered node in the graph G and j is the j^th^ entry in query Y.

p^i,m^: predecessor node which is m edges back of the i^th^ topologically ordered node in graph G.

Fig.1 shows an example where the equation is applied to each cell in the matrix, starting from the top-left cell, dp(0, 0), where i = 0 and j = 0. The alignment scores for each cell are computed row by row until the bottom-right cell is reached. At dp(3, 2), the first 2-mer (“AC”) is encountered, and the score is updated using dp(3, 2) = dp(1, 1) + 2 = 2. For the final cell in the matrix, the score is calculated as dp(7, 4) = dp(5, 2) + 2 = dp(2, 1) + 4 = 4.

**Fig. 1.**
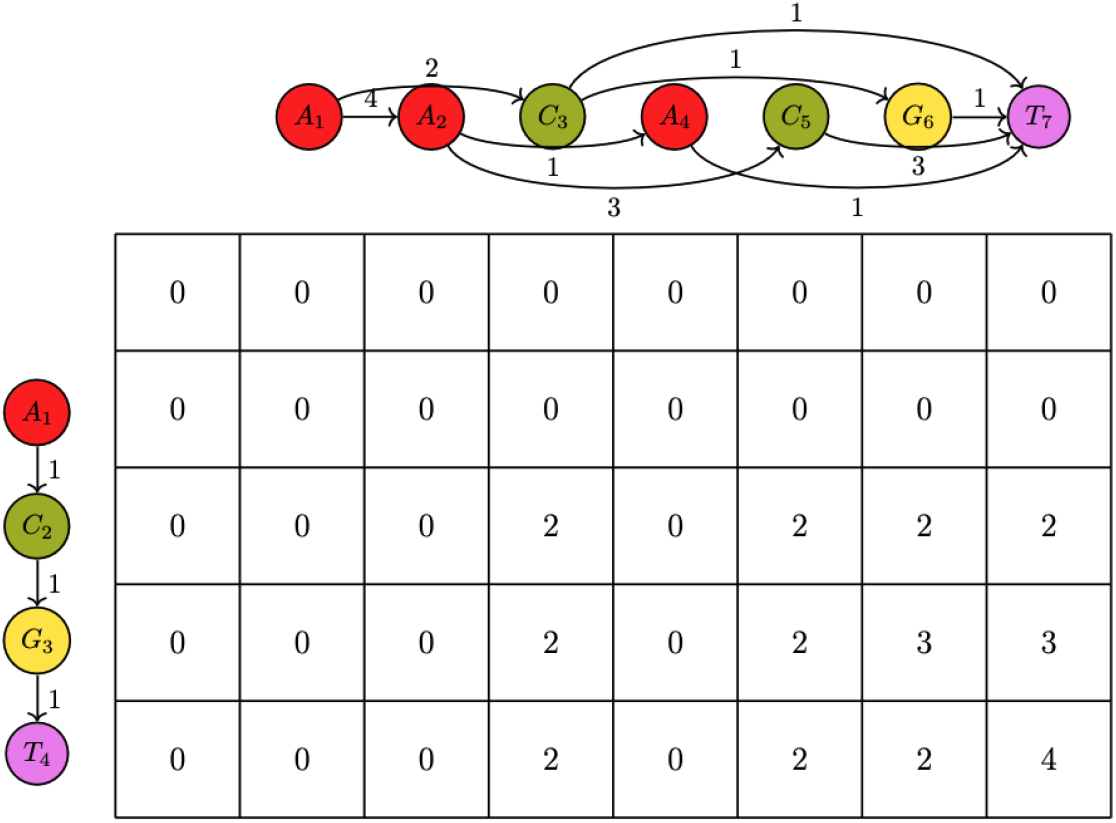
Basic DP programming example, k = 2, each cell show dp(i, j) which is the LCSk++ alignment score for i^th^ row j^th^ column

The time complexity of this algorithm is O(LM * min(L, M)) where L is query length and M is number of nodes in graph G. Performing basic dynamic programming and obtaining the LCSk++ path for banded POA is not a viable solution, as the time complexity of basic dynamic programming for LCSk++ alone is higher than that of full POA.

#### Efficient algorithm

An efficient algorithm presented by [10] achieves a significant speed-up from the basic dynamic programming algorithm and is a crucial component in banded sequence-to-sequence dynamic programming alignment[13]. We propose a query-to-graph version that could achieve similar improvements in partial order alignment. The definition of a match pair is described as follows to accommodate a graph structure, modified from the original definition [12]. Definitions:

s^i, m^ = successor node which is m edges in front of the i^th^ topologically ordered node in graph G.

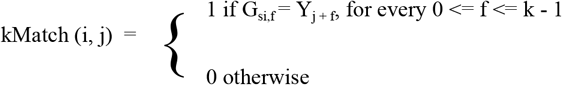

If kMatch(i, j) = 1, we call (i, j) a match pair. (i, j) is called the start of the match pair and (s^i,k-1^, j + k - 1) is called the end of the match pair, this in contrast to the end match pair of the original definition as there could be multiple ends if (s^i,k^, j + k) is used. We adjust the algorithm to incorporate this change. (Fig. 1) shows an example of the efficient algorithm.

The formula for the efficient algorithm is as follows,

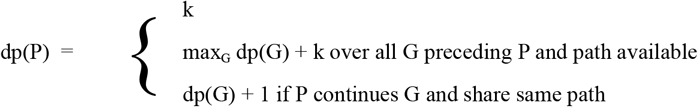

kmer match pair P can start its k-common subspace, or extend a k-common subspace if there is a path available from start of P to end of G or extend a k-common subsequence by enlarging k if the continuing nodes are on the same path.

In the example shown in Fig. 2, once the start and end positions of the k-mers are identified, the equation dp(P) is applied to each end position. The computation proceeds row by row, starting from the top-right corner. The end of c is reached first, where dp(c-end) = k = 2, as no preceding k-mer exists. Next, the end of a is processed, yielding dp(a-end) = k = 2. The end of d follows, with dp(d-end) = k = 2, as no prior k-mer is found. Subsequently, the end of e is reached, where dp(e-end) = dp(c-end) + k = 2 + 2 = 4, (as c precedes e). Finally, the end of b is processed, with dp(b-end) = dp(a-end) + 1 = 3, (as b continues a). Thus, the LCSk++ path is identified as [e-end, c-end], containing two k-mers with a total score of 4.

**Fig. 2.**
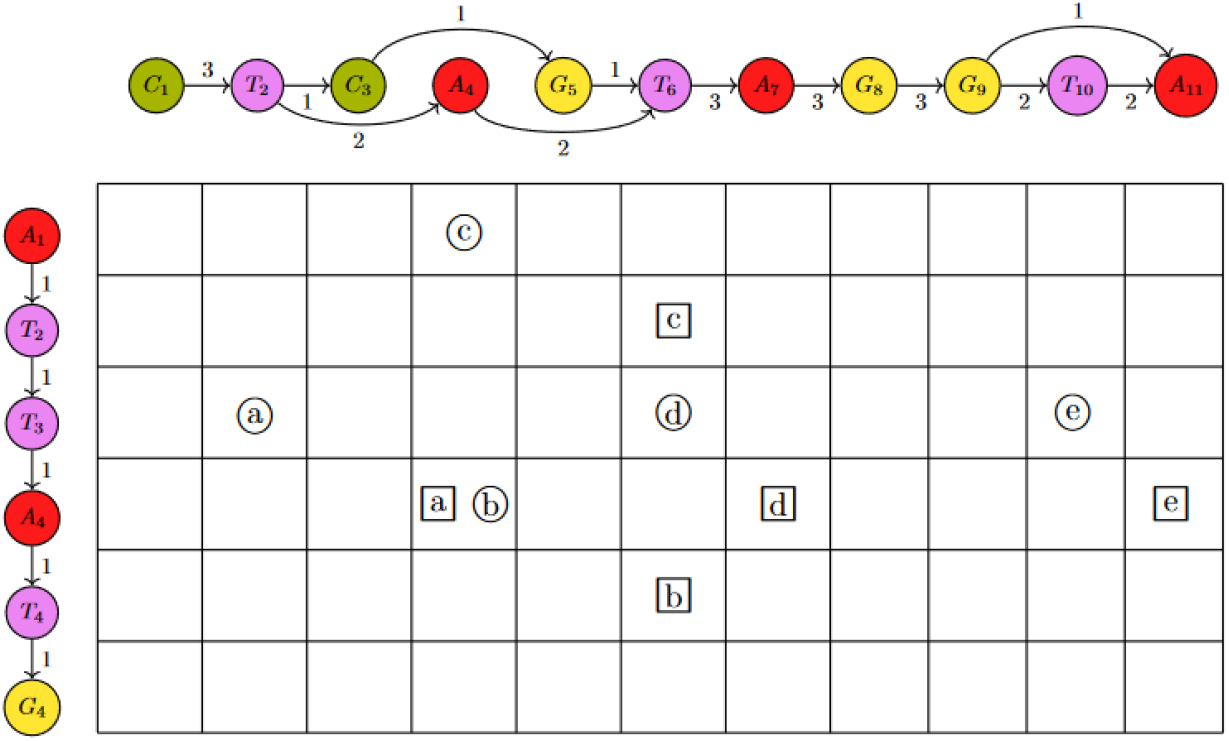
Matching kmers in query to POA graph, k = 2; 5 kmer match pairs are present denoted a to e. Starts are presented by the circles and ends by squares. “b continues a”, “c precedes e” applies.

#### Efficient algorithm implementation

##### Algorithm 1

Sequence to graph LCSK++ computation

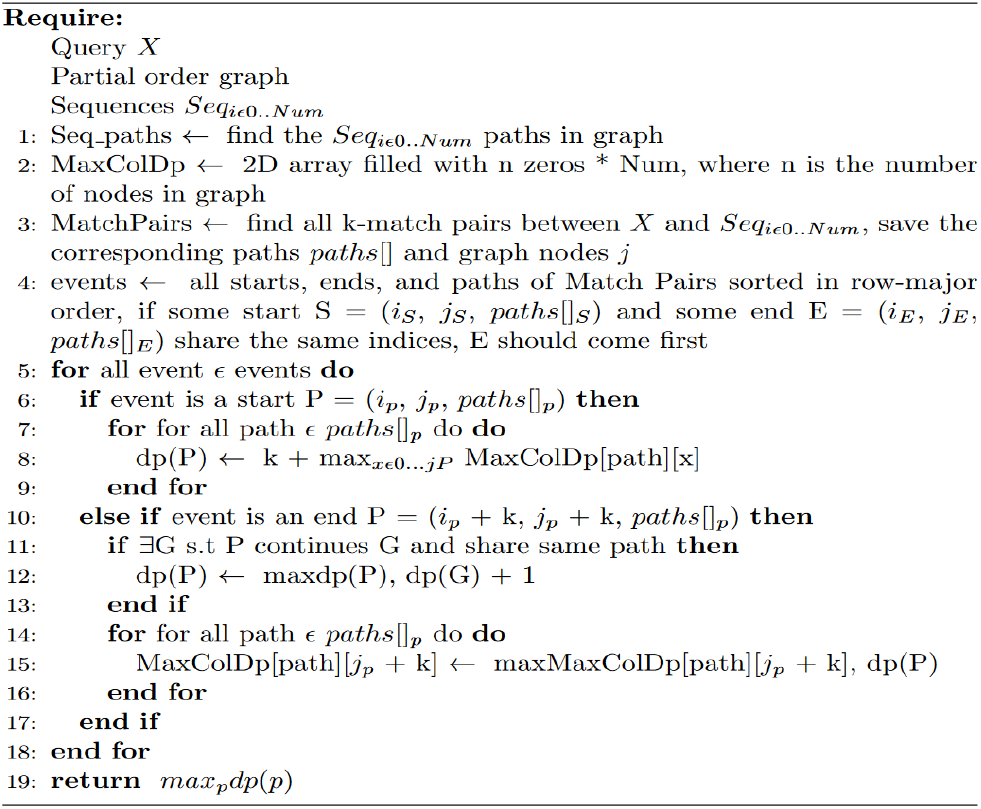

The following variables and notations are used throughout the algorithm and the time complexity calculation:

V: vertices in graph

E: edges in graph

Num: number of sequences

n: query length

m: max sequence length

r: number of kmer matches between query and sequences

The algorithm obtains the max kmer path of a query sequence X to an existing partial order alignment (POA) graph using LCSkPOA algorithm. The process is illustrated in Fig. 2, which provides a step-by-step example. The inputs to the algorithm include the query sequence X, the current POA graph (comprising the previously aligned sequences), and the sequences aligned to the POA graph. The output is the LCSk++ path with max kmers. The algorithm proceeds as follows:

Line 1: Using a depth-first search (DFS) algorithm, the sequence paths in the POA graph are identified. The DFS leverages the previously aligned sequences (Num) to ensure all relevant paths are accounted for.

Line 2: A 2D array of dimensions Num×V is initialized. This array will store the maximum column values for each node in the graph, corresponding to each sequence path.

Line 3: All k-mer match pairs between the query sequence X and the graph paths are identified. This step involves hashing the query sequence and identifying corresponding matches in each graph path.

Line 4: The k-mer matches are sorted in row-major order. For each match, the algorithm records relevant details such as start and end positions, paths, and the corresponding graph nodes. These details are encapsulated in an event structure, which serves as the basis for subsequent processing.

Line 5: The algorithm processes events one by one to update alignment scores and column values.

Lines 6–7: If the event represents the start event, the d(P) value for the current path is updated. The value is computed as the maximum column value for the current path + k.

Lines 11–13: If the event continues from a previous event, the d(P) value is updated using the formula d(P) = d(G) + 1, where G represents the predecessor.

Lines 14–16: After processing an event, the maximum column values for all paths are updated in the 2D array.

Line 19: The MaxColDP 2D array is returned, this includes the max kmer path.

The complexity of various operations can be analyzed as follows. Finding sequence paths in the graph has a time complexity of O(Num * (V + E)). Finding match pairs involves hashing the query in O(n) and matching in O(Num * m), resulting in O(n + Num * m). Sorting events has a complexity of O(r * log(r)). Lines 7 to 15 are handled by Num Fenwick trees, which operate in O(Num * log n). Processing all events takes O(r * Num * log n). Consequently, the total time complexity is O(Num * (V + E) + n + Num * m+ r log r + r * Num * log n), and the total space complexity is O(r). By integrating sequence finding in the graph using DFS while recording nodes during the POA, the time complexity can be reduced to O(n + Num * m + r log r + r * Num * log n), significantly enhancing performance when aligning a large number of sequences in the POA graph.

### Using LCSk++ backbone in POA

After obtaining the LCSk++ path, which is the path with the most k-mers, we need an optimal method to use this path to create the bands for the POA dynamic programming matrix. Initially, we band the entire section until the first k-mer is encountered. Similarly, we band from the last k-mer to the end, as illustrated by the yellow dot pattern region in (Fig. 3). This ensures the region necessary for semi-global and local alignment is present. Second, for the k-mer matches, we create a specified band around them, indicated by the orange vertical line pattern region in (Fig. 3). Third, in the intersections between k-mer matches, we band the entire area, as shown in the purple horizontal line pattern region in (Fig. 3). The band size is chosen based on the sequence length; for most PacBio datasets, a band size of 200 has proven effective.

**Fig. 3.**
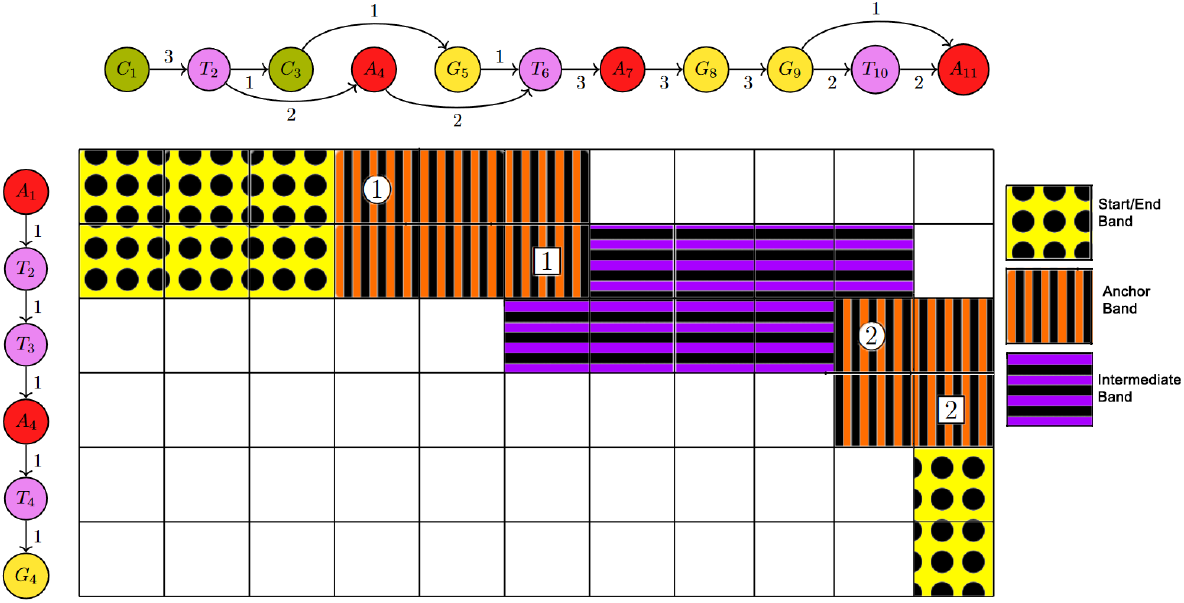
Banding using the LCSk++ path: Yellow dot pattern represents the start and end banding which enables semi-global/local alignment, orange vertical line pattern indicates the LCSk++ path anchor banding (1 and 2 align with *c* and *e* in Fig. 2 as the max 2-mer path; circles and squares represent their starts and ends, respectively), and purple horizontal pattern represents the intermediate section banding between the anchors.

### Parallel Processing

This section demonstrates how parallel processing can be implemented following the identification of the LCSk++ path by partitioning the POA matrix into independent sections.

The result after obtaining the LCSk++ path is shown in Fig. 4, where the identified k-mers serve as candidates for partitioning points. The graph nodes and query are traversed to evaluate candidates for partitioning. A predefined upper limit is imposed on the number of nodes per partition and the query bases. Once this limit is reached, potential partitioning candidates are assessed. Specifically, if a candidate node in the POA graph has only a single incoming edge with a weight corresponding to the number of sequences aligned, it can be designated as a partitioning point, as illustrated in Fig. 5B.

**Fig. 4.**
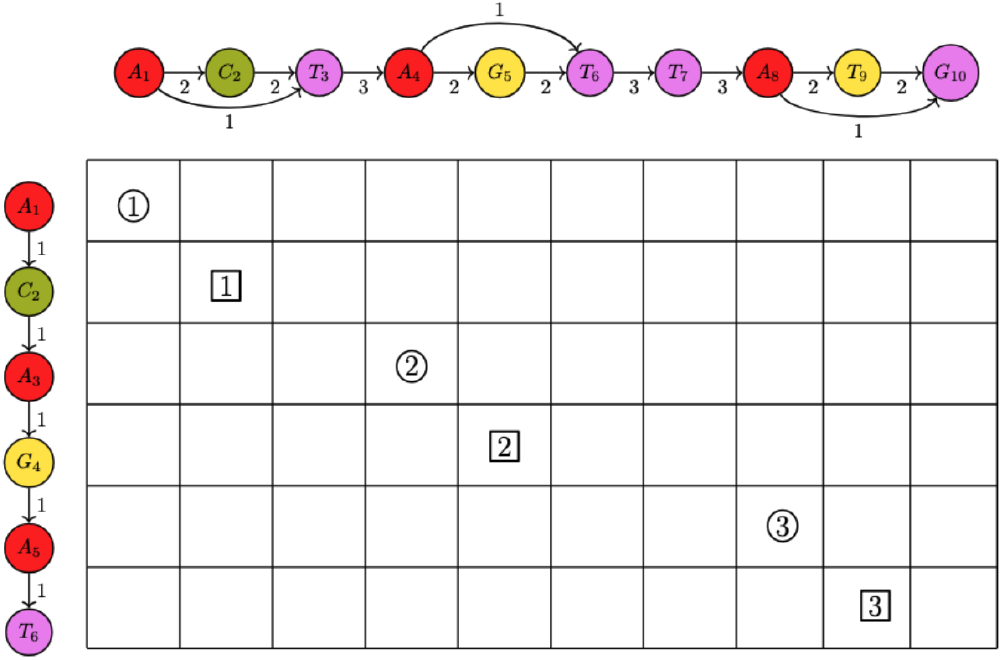
LCSk++ path identified, with potential candidates for partitioning points indicated. Anchors 1, 2, and 3 represent 2-mers on the maximum k-mer path, with their starts denoted by circles and ends by squares. These anchors are evaluated as candidates for partitioning based on the POA graph structure.

**Fig. 5.**
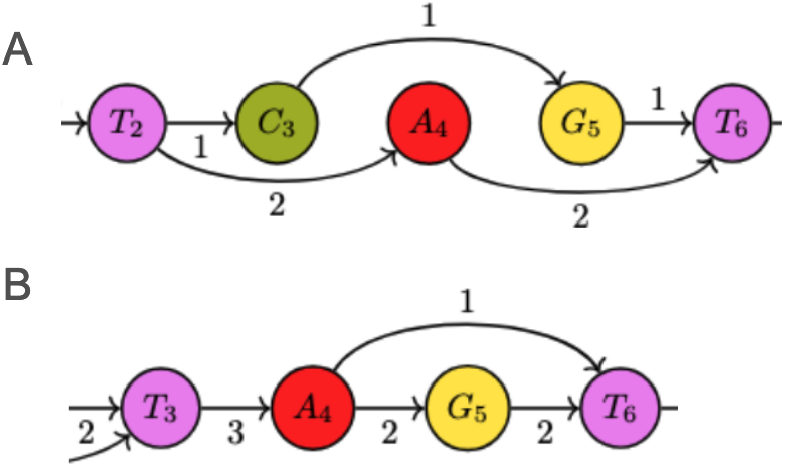
A. Candidate node A_4_ of POA graph not suitable for partitioning, B. Candidate node A_4_ of POA graph suitable for partitioning.

This process is applied recursively until the end of the graph and query sequence is reached. The identified partitioning points allow independent sections of the matrix to be processed in parallel. Separate threads can be employed for each of these partitions, with the main thread subsequently concatenating the results to generate the final POA alignment for the complete matrix.

The parallel processing approach uses less memory than single-threaded processing because the cells around the partition points in the multithreaded method are not fully banded. This reduction is possible due to optimal partition point selection, which ensures that the backtrace passes through these points. This difference is illustrated in Fig. 6. In the multithreaded approach (Fig. 6A), only 20 matrix cells are computed, whereas the single-threaded approach (Fig. 6B) requires 24 cells. This method leverages modern computational resources to achieve significant speed improvements and reduced memory consumption, while maintaining high alignment quality due to optimal partition point selection. It is particularly advantageous for processing large-scale genomic datasets.

**Fig. 6.**
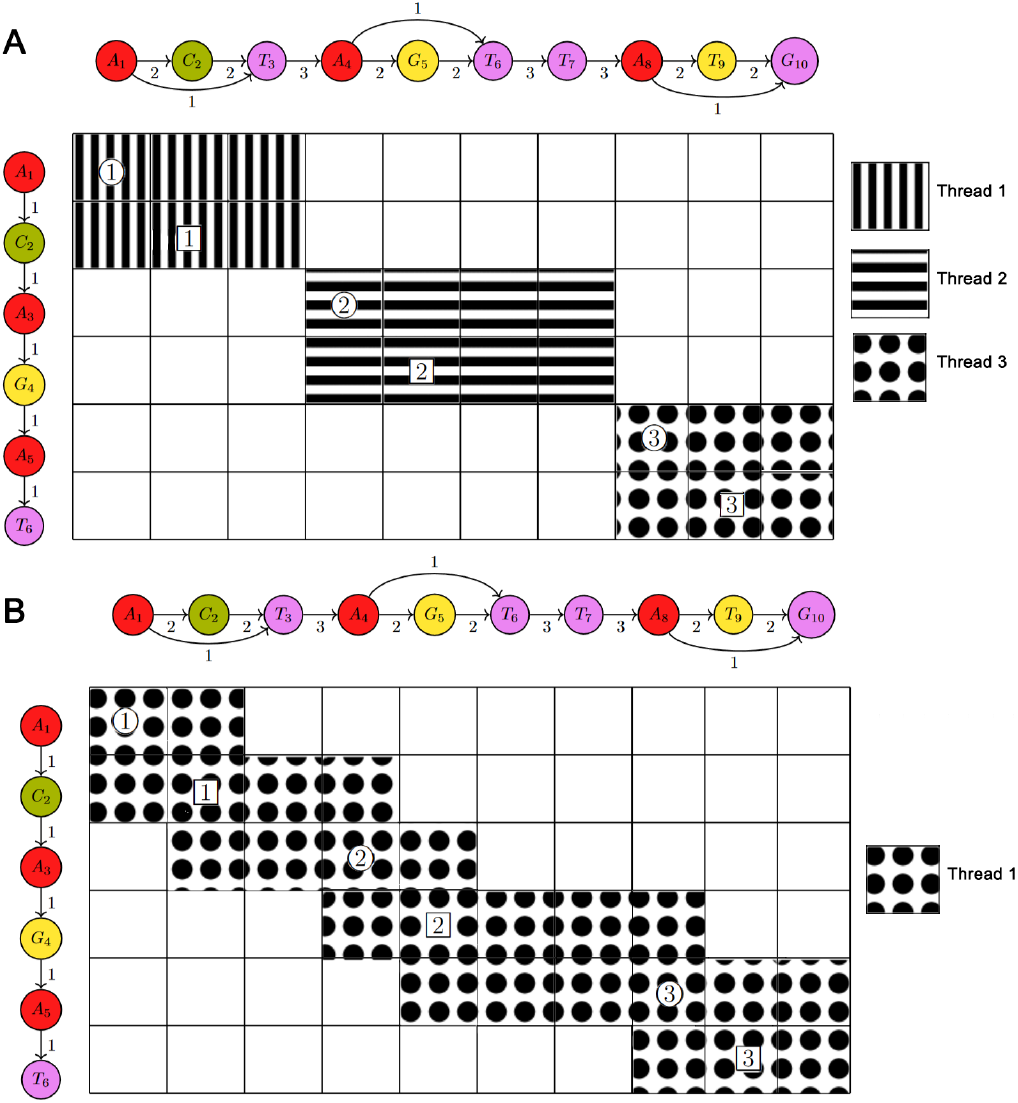
A. Multithreaded banded POA matrix. B. Single-threaded banded POA matrix. In both, 1, 2, and 3 represent 2-mers from max 2-mer path, with starts denoted by circles and ends by squares.

**Fig. 7.**
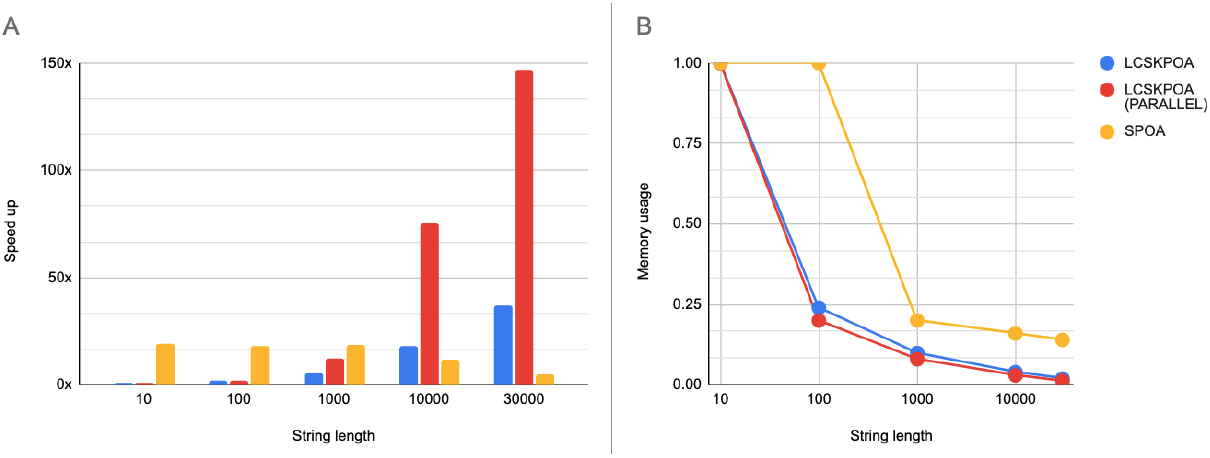
Resulting graphs of running LCSkPOA with Synthetic data. A.speed up of SPOA, LCSkPOA and LCSkPOA parallel compared with vanilla semi-global POA B. Memory usage of SPOA, LCSkPOA and LCSkPOA parallel compared with vanilla semi-global POA.

**Fig. 8.**
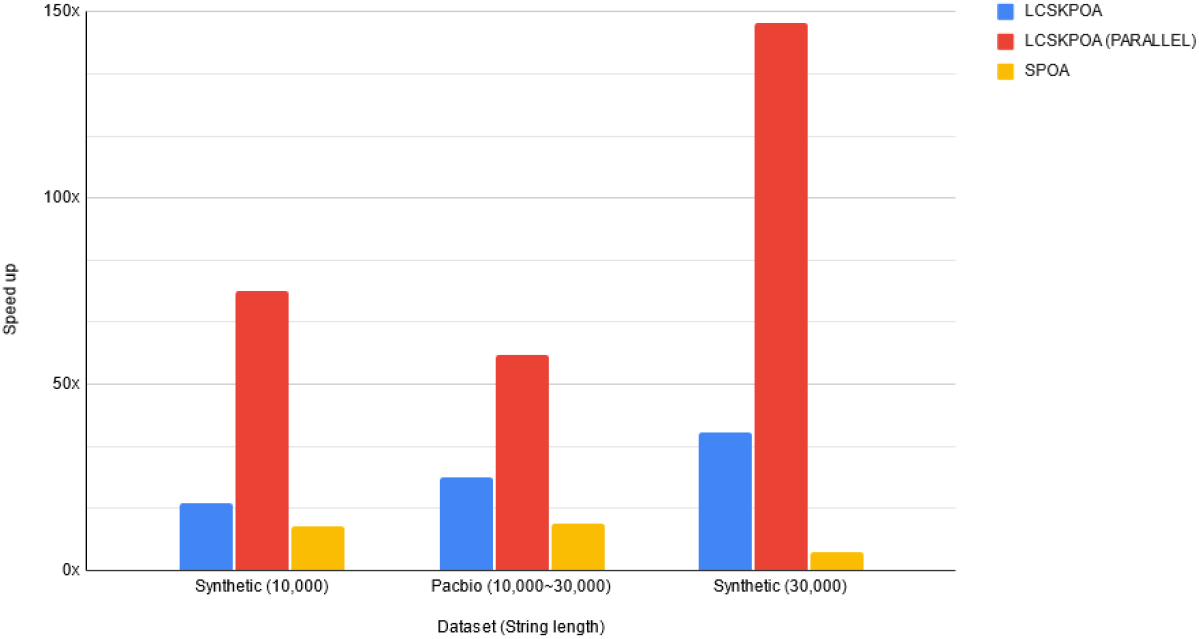
Comparison of Speed up of LCSkPOA and LCSkPOA parallel when run with synthetic dataset and PacBio HiFi data set.

## Results

### System configuration

CPU: AMD EPYC 7313 16-Core Processor 64 threads

Memory: DDR4 420GB

Language used: Rust 1.65.0

Libraries used: Rust-bio for POA, std::thread for parallel processing implementation

### Synthetic data result

The synthetic data set was generated by the custom sequence generator for this project. The custom code introduces variations like insertions, deletions, and substitutions, simulating realistic genetic changes for testing our methods. The PacBio dataset used in this study was obtained from real sequenced data, the same as that utilized in topoqual [14].

(Fig. 3A) shows the speed-up factors for LCSkPOA, LCSkPOA (PARALLEL), and SPOA compared to vanilla unbanded POA across different string lengths. LCSkPOA (PARALLEL) achieves dramatic speed increases, reaching up to 150x for the longest strings, while LCSkPOA shows modest improvements. SPOA achieves high speed-up for short sequences but remains low for long sequences. (Fig. 3B) illustrates memory usage for the same methods, with LCSkPOA and its parallel version displaying steep declines in memory consumption for longer strings and stabilizing at low levels. In contrast, SPOA initially uses high memory but reduces significantly for longer strings due to architectural and language optimizations. However, LCSkPOA (PARALLEL) still maintains memory usage that is tenfold lower than SPOA for a string length of 30,000. These results highlight the efficiency of the LCSkPOA algorithm, especially with parallel processing for longer sequences, in providing substantial speed-up and maintaining low memory usage, making it a robust solution for local and semi-global alignment in Partial Order Alignment (POA) graphs.

### Pacbio data vs synthetic data result

In (Fig. 4), the single-threaded LCSkPOA exhibits a linear speedup from synthetic datasets of length 10,000 to PacBio HiFi datasets of length 10,000 to 30,000 and to synthetic datasets of length 30,000. However, the multi-threaded LCSkPOA does not demonstrate linear scaling for the PacBio HiFi datasets which are around 10,000 ∼ 30,000 in length. This discrepancy is due to the presence of edges that span the majority of the graph in PacBio HiFi datasets as a result of the multiple repeating sections in the human genome, making it more challenging to find suitable anchor points for partitioning. As a result, parallel processing is less effective for PacBio HiFi datasets compared to synthetic datasets. SPOA performance becomes weaker as the string lengths increase.

**Table 1.** Comparison of run time, memory usage, and accuracy with vanilla semi-global poa with LCSkPOA with different K values (Averages of 20 runs, Synthetic data and PacBio HiFi data)

|  | Unbanded POA |  |  | LCSkPOA (Parallel) |  |  |  |
| --- | --- | --- | --- | --- | --- | --- | --- |
| String length | Score | Run time | Memory | k | POA Alignment Score | Run time | Memory |
| **10** | 16 | 4μs | 8KB | 2 | 16<br>(100%) | 21μs<br>(100%+) | 4KB<br>(50.8%) |
| | | | | 4 | 16<br>(100%) | 14μs<br>(100%+) | 4KB<br>(50.8%) |
| **100** | 142 | 458μs | 462KB | 4 | 142<br>(100%) | 499.85μs<br>(100%+) | 86KB<br>(18.7%) |
| | | | | 8 | 128<br>(98.6%) | 326μs<br>(71.2%) | 107KB<br>(23.2%) |
| **1,000** | 1,387 | 46ms | 40MB | 4 | 1,300<br>(100%) | 4.3ms<br>(9.3%) | 8.7MB<br>(21.7%) |
|  |  |  |  | 8 | 1,362<br>(99.1%) | 3.7ms<br>(8.1%) | 2.5MB<br>(6.25%) |
| **10,000** | 13,944 | 4.6s | 5.3GB | 8 | 13,945<br>(100%) | 807.02ms<br>(17.5%) | 182MB<br>(3.4%) |
|  |  |  |  | 12 | 13,820<br>(99.1%) | 64.5ms<br>(1.4%) | 230MB<br>(4.3%) |
| **30,000** | 41,715 | 41.3s | 55GB | 8 | 41,715<br>(100%) | 1,720ms<br>(4.1%) | 827MB<br>(1.55%) |
|  |  |  |  | 12 | 41,351<br>(99.1%) | 301ms<br>(0.7%) | 694MB<br>(1.25%) |
| **10,000~30,000**<br>**(PacBio HiFi)** | 29,165 | 17s | 18.8GB | 8 | 29,458<br>(100%) | 3,960ms<br>(23.2%) | 677MB<br>(3.76%) |
|  |  |  |  | 12 | 29,458<br>(100%) | 400ms<br>(2.3%) | 299MB<br>(1.58%) |

The table provides a comparative analysis of vanilla unbanded semi-global POA and LCSkPOA (Parallel) across different string lengths. It highlights the influence of the parameter *k* on POA alignment scores—calculated with a scoring scheme of +2 for matches and -2 for mismatches and gaps—as well as on runtime and memory usage. For short string lengths, such as 10 and 100, vanilla POA achieves high POA alignment scores with low run times and minimal memory usage. In comparison, LCSkPOA (Parallel) also achieves high POA alignment scores, but with slightly higher run times and significantly lower memory usage. As the string length increases to 1,000, vanilla POA’s run time and memory usage grow substantially, whereas LCSkPOA (Parallel) maintains high POA alignment scores and demonstrates significant efficiency improvements. Notably, as k increases from 4 to 8, the run time decreases from 4.3ms to 3.7ms, indicating no significant gain in using higher k values for short and medium-length sequences.

For longer string lengths such as 10,000 and 30,000, the benefits of LCSkPOA (Parallel) become more evident. vanilla POA exhibits a drastic increase in both run time and memory usage, reaching 4.6s and 5.3GB for 10,000 strings, and 41.3s and 55GB for 30,000 strings. Conversely, LCSkPOA (Parallel) with higher k values significantly reduces run time and memory usage. For example, at a string length of 10,000 and k=12, the run time is 64.5ms, reduced from 807ms at k=8, while maintaining a high score of 99.1%. At 30,000 strings and k=12, LCSkPOA (Parallel) achieves a run time of 301ms, reduced from 1,720ms at k=8, showcasing a stark contrast to vanilla POA.

For a real sequencing dataset, PacBio HiFi (10,000∼30,000), LCSkPOA (Parallel) with k=12 achieves a run time of 400ms, reduced from 3,960ms at k=8, maintaining a perfect score of 100%. This comparison clearly highlights the significant impact of increasing k in LCSkPOA (Parallel) for long sequences, consistently improving run time and memory efficiency while maintaining near perfect POA alignment scores, making it highly effective for large-scale sequence alignment tasks.

## Conclusion

In this study, we introduced an innovative adaptation of the Longest Common Subsequence with k-mer matches (LCSk++) algorithm, tailored specifically for graph structures and particularly for Partial Order Alignment (POA) graphs. POA graphs, which are directed acyclic graphs, effectively represent multiple sequence alignments and capture the relationships between sequences with more flexibility than traditional alignment methods, accommodating variations such as insertions, deletions, and substitutions.

Our approach extends the LCSk++ algorithm, originally designed for linear string sequences, to manage the complexities of graph data structures by integrating dynamic programming and graph traversal techniques. This extension allows for the detection of conserved regions within POA graphs, referred to as the LCSk++ backbone,

which facilitates banding of the POA matrix for accurate local and semi-global alignment. This method significantly improves the construction of consensus sequences with high accuracy and efficiency and supports other critical applications of POA, such as multiple sequence alignment (MSA) for phylogeny construction and graph-based reference alignment.

The results of our study demonstrate a substantial enhancement in performance, with a 37-fold increase in speed and a 98% reduction in memory usage for sufficiently long sequences. This reduction in memory usage is particularly beneficial, allowing up to 150x speed improvements on conventional PCs through parallel processing without compromising alignment quality, a feat not achievable with vanilla unbanded POA. These advancements enable fast and efficient local and semi-global alignment on the banded POA graph, significantly enhancing its utility in bioinformatics.

Overall, the extended LCSk++ algorithm for graph structures represents a significant advancement in sequence alignment technology, offering improved performance and accuracy for the analysis of complex biological datasets, thereby addressing some of the longstanding challenges in multiple sequence alignment.

## Availability and requirements

**Project name:** lcskgraph

**Project home page:** https://github.com/lorewar2/lcskgraph

**Operating system(s):** Linux, MacOS

**Programming language:** Rust

**Other requirements:** Rust 1.65.0+

**License** MIT license

**Any restrictions to use by non-academics:** None

### Abbreviations

POA: partial order alignment
LCSk: Longest Common Subsequence K-mer
SIMD: Single Instruction, Multiple Data

## Declarations

## Ethics approval and consent to participate

We acquired a cord blood sample (ID: PD47269d) from a newborn female patient through the Cambridge Blood and Stem Cell Biobank (CBSB). This sample was collected at Addenbrooke’s Hospital with informed consent and approved by the Cambridge East Ethics Committee under reference 18/EE/0199.

## Consent for publication

Not applicable

## Availability of data and materials

The datasets analysed during the current study are available in the github repository, https://github.com/lorewar2/lcskgraph and can also be accessed by SRA repository, SRA biosample accession PRJNA1128051.

## Authors’ contributions

Methodology, MW, HH; Conceptualization, MW, CS; Resources, HH; Supervision, HH, CS; Writing original draft, MW, CS, HH; Validation, MW;

All authors reviewed the manuscript. All authors read and approved the final manuscript.

## Acknowledgements

Not applicable

